# Antimalarials in mosquitoes overcome *Anopheles* and *Plasmodium* resistance to malaria control strategies

**DOI:** 10.1101/2021.03.12.435188

**Authors:** Douglas G. Paton, Alexandra S. Probst, Erica Ma, Kelsey L. Adams, W. Robert Shaw, Naresh Singh, Selina Bopp, Sarah K. Volkman, Domombele F. S. Hien, Prislaure S. L. Paré, Rakiswendé S. Yerbanga, Abdoullaye Diabaté, Roch K. Dabiré, Thierry Lefèvre, Dyann F. Wirth, Flaminia Catteruccia

**Affiliations:** Department of Immunology and Infectious Diseases, Harvard TH Chan School of Public Health, Boston, USA.; Institut de Recherche en Sciences de la Santé/Centre Muraz, Bobo-Dioulasso, Burkina Faso; MIVEGEC, IRD, CNRS, University of Montpellier, Montpellier, France; Laboratoire mixte international sur les vecteurs (LAMIVECT), Bobo Dioulasso, Burkina Faso; Centre de Recherche en Écologie et Évolution de la Santé (CREES), Montpellier, France

## Abstract

The spread of insecticide resistance in *Anopheles* mosquitoes and drug resistance in *Plasmodium* parasites is contributing to a global resurgence of malaria, making the generation of control tools that can overcome these issues an urgent public health priority. We recently showed that the transmission of *Plasmodium falciparum* parasites can be efficiently blocked when exposing *Anopheles gambiae* females to antimalarials deposited on a treated surface, with no negative consequences on mosquito fitness. Here, we demonstrate this approach can overcome the hurdles of insecticide resistance in mosquitoes and drug resistant in parasites. We show that the transmission-blocking efficacy of mosquito-targeted antimalarials is maintained when field-derived, insecticide resistant *Anopheles* are exposed to the potent cytochrome b inhibitor atovaquone, demonstrating that this drug escapes insecticide resistance mechanisms that could potentially interfere with its function. Moreover, this approach prevents transmission of field-derived, artemisinin resistant *P. falciparum* parasites (*Kelch13* C580Y mutant), proving that this strategy could be used to prevent the spread of parasite mutations that induce resistance to front-line antimalarials. Atovaquone is also highly effective at limiting parasite development when ingested by mosquitoes in sugar solutions, including in ongoing infections. These data support the use of mosquito-targeted antimalarials as a promising tool to complement and extend the efficacy of current malaria control interventions.

**Significance Statement:** Effective control of malaria is hampered by resistance to vector-targeted insecticides and parasite-targeted drugs. This situation is exacerbated by a critical lack of chemical diversity in both interventions and, as such, new interventions are badly needed. Recent laboratory studies have shown that an alternative approach based on treating *Anopheles* mosquitoes directly with antimalarial compounds can render the vector incapable of transmitting the *Plasmodium* parasites that cause malaria. While promising, showing that mosquito-targeted antimalarials remain effective against wild parasites and mosquitoes, including drug- and insecticide-resistant populations, respectively, is crucial to the future viability of this approach. In this study, carried out in the US and Burkina Faso, we show that antimalarial exposure is highly effective, even against extremely resistant mosquitoes, and can block transmission of drug-resistant parasites. By combining lab, and field-based studies in this way we have demonstrated that this novel approach can be effective in areas where conventional control measures are no longer as effective.

## Introduction

Human malaria, a parasitic disease caused by unicellular eukaryotic *Plasmodium* parasites and spread through the bite of *Anopheles* mosquitoes, remains a substantial cause of global morbidity and mortality (1). Malaria control programs rely on preventative measures focused on mosquito control and on therapeutic measures based on the use of antimalarial drugs. Mosquito-targeted interventions are the most effective tools at reducing the transmission of *Plasmodium* parasites, with long-lasting insecticide-impregnated nets (LLINs) and indoor residuals spraying (IRS) as primary methods for malaria prevention. LLINs alone are predicted to have contributed to 68% of malaria cases averted between 2000 and 2015 (2). Alongside these preventative interventions, artemisinin combination therapies (ACT) have been the cornerstone of human malaria treatment since their widespread introduction at the beginning of this century (3, 4) and have contributed substantially to the reduction in malaria mortality and morbidity observed since then (2).

Despite sizeable investment, malaria control and elimination efforts are, however, faltering due to reduced operational effectiveness of these key control tools, largely caused by mosquito resistance to insecticides and parasite resistance to drugs (5–8). In the malaria hyperendemic regions of southern Mali and southwest Burkina Faso, for example, resistance to pyrethroids is extreme (9), driven by multifactorial and synergistic resistance mechanisms including enhanced metabolic detoxification through upregulated cytochrome P450s (metabolic resistance), and reduced tarsal uptake through cuticular thickening (cuticular resistance) (10–13). Similarly, the emergence and spread of artemisinin resistance to sub-Saharan Africa — a region where as many as 93% of annual malaria deaths occur (1) — is a major concern. Until recently, resistance to these first line antimalarials was limited geographically to the Greater Mekong Subregion (GMS), however *de novo* mutations in *PfKelch13* associated with *in vitro* resistance have now been detected in Uganda, Tanzania, and Rwanda (8, 14–19). Of further concern is the recent invasion and spread of the Asian vector *Anopheles stephensi* to the horn of Africa (20), as this mosquito species is highly competent for the transmission of *P. falciparum* parasites endemic to the GMS and therefore, invasive populations may facilitate the spread of parasites harboring artemisinin resistance mutations from Asia to Africa. Besides insecticide resistance, an additional hurdle to malaria elimination is represented by residual malaria. Defined as malaria transmission in the presence of universal effective LLIN coverage, residual malaria is driven by mosquitoes that exhibit outdoor or daytime biting preferences and is a considerable hurdle to malaria eradication efforts (21). Increased focus on residual malaria has stimulated interest in the use of attractive toxic, or targeted (22), sugar baits (ATSBs) to attract and kill adult mosquitoes irrespective of blood-feeding behavior, which in field trials have shown some promise as a tool for suppressing vector populations (23, 24).

Thus, a control strategy that prevents insecticide resistant *Anopheles* populations from transmitting malaria parasites, including parasite strains carrying drug resistance mutations, regardless of mosquito feeding behavior and without imposing strong selective pressure on mosquitoes could overcome the limitations of current mosquito-targeted interventions. In an effort to generate such a strategy, we recently demonstrated that transmission of *P. falciparum* parasites can be prevented when *Anopheles gambiae* are exposed to the antimalarial atovaquone (ATQ) through direct contact — analogous to the mode of insecticide exposure on LLINs or IRS (25). Contact with ATQ-coated surfaces completely abrogated parasite development when it occurred around the time of infection (between 24 hours (h) before and 12 h post feeding on *P. falciparum*-infected blood), preventing onward transmission of the parasite. Importantly, mathematical models based on these results showed that integrating antimalarial ingredients into existing mosquito-targeted interventions could considerably reduce malaria transmission in areas of widespread insecticide resistance, empowering our best malaria prevention tools (25).

Here we show that targeting *P. falciparum* with antimalarials during its development in the *Anopheles* female circumvents the hurdles of insecticide and drug resistance, providing a critical addition to the malaria elimination toolkit. Parasite development is substantially reduced when wild, as well as recently lab adapted *Anopheles coluzzii* (a sibling species of *An. gambiae*) that are highly resistant to pyrethroids are exposed to ATQ prior to feeding on blood taken from *P. falciparum-*infected donors in Burkina Faso. ATQ is also fully active against field-derived *P. falciparum* parasites from Cambodia that are resistant to artemisinin. When using distinct drug targets in humans and mosquitoes, this method is therefore capable of both overcoming insecticide resistance mechanisms and stopping transmission of parasite mutations that confer resistance to frontline antimalarials. Finally, we show that delivering ATQ via sugar solutions causes a striking reduction in both parasite numbers and growth, proving that antimalarials could also be incorporated into interventions that target outdoor malaria transmission, such as ATSB. Targeting *Plasmodium* parasites in the mosquito vector is therefore a promising strategy that circumvents key limitations of current malaria control and preventative interventions.

## Results

### Exposure to ATQ substantially reduces infection with field *P. falciparum* isolates in insecticide resistant *An. coluzzii*

To determine whether antimalarial exposure can maintain efficacy in insecticide-resistant mosquitoes, we took *An. coluzzii* collected as pupae from larval breeding sites in Bama, Burkina Faso and reared them to adults at the IRSS, Burkina Faso. Adult mosquitoes were infected using *P. falciparum* gametocyte positive blood obtained from a malaria infected human donor on the day of infection. The *An. coluzzii* mosquitoes endemic to this part of Burkina Faso – hereafter named AcVK5 – are highly resistant to pyrethroids (9, 11, 26). AcVK5 mosquitoes were exposed to ATQ for 6 minutes (min) at two concentrations (100 µmol- or 1 mmol/m^2^) or a mock-treated blank control surface prior to feeding on infectious blood samples. Infection outcomes were assayed at 7 days (d) post infectious blood meal (pIBM) by dissection of the mosquito midgut to determine the prevalence and intensity of parasite oocysts (Fig. 1 a). Control-exposed AcVK5 females were robustly infected with 81.3% harboring at least one *P. falciparum* oocyst, and median infection intensity in infected females of 19 oocysts per midgut. In contrast, AcVK5 mosquitoes exposed to either dose of ATQ had significantly reduced *P. falciparum* infection both in terms of prevalence and intensity (Fig 1b). At the highest concentration, we observed a 99% overall reduction in infection relative to the control (84.6% reduction in prevalence and 94.8% reduction in median intensity), while at the lower dose inhibition of infection reached 96% overall (65.9% reduction in prevalence and 89.5% reduction in median intensity). These results demonstrate that direct tarsal antimalarial exposure, for instance incorporating antimalarials in LLINs and IRS, can effectively block transmission of circulating west African *P. falciparum* parasites in highly insecticide resistant endemic *Anopheles* mosquitoes. Although our previous findings had shown that both ATQ doses tested above are capable of complete inhibition of parasite transmission using the combination of standard *P. falciparum* (NF54) and insecticide-susceptible *Anopheles* (G3) populations (25), our results confirm that parasite development can be considerably impaired when exposing mosquitoes to antimalarials.

**Figure 1:**
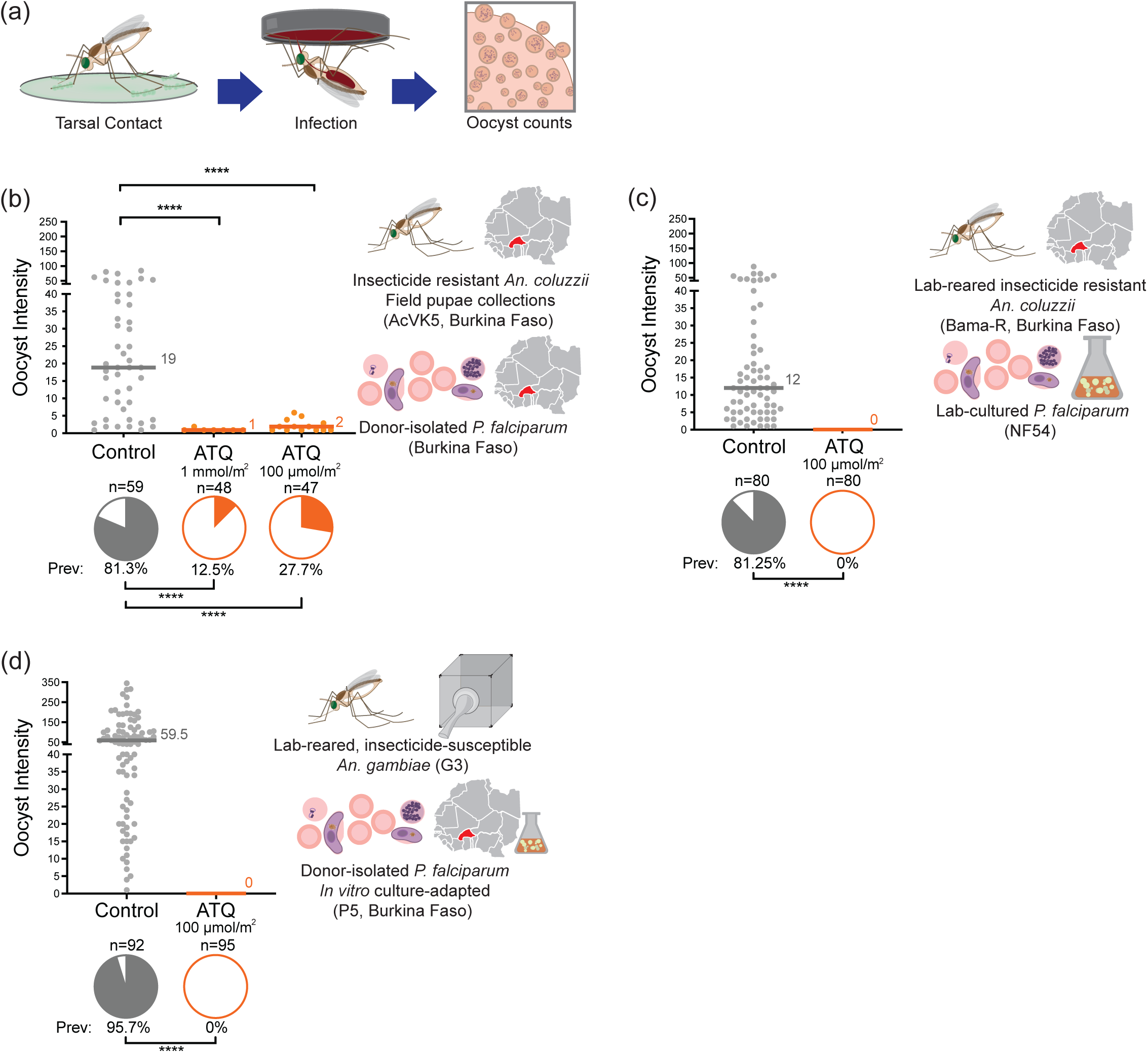
Exposing *An. gambiae* mosquitoes to ATQ suppresses *P. falciparum* development in field-derived parasites and insecticide-resistant mosquitoes. (**a**) Experimental scheme. (**b**) Female AcVK5 mosquitoes, collected as pupae from breeding sites in southwestern Burkina Faso, were exposed to ATQ by tarsal contact at either the maximal effective concentration (EC_99_) for insecticide-susceptible mosquitoes (100 µmol/m^2^, EC_99_) or 10x EC_99_ (1 mmol/m^2^) for 6 minutes, prior to infection with donor-isolated “wild” *P. falciparum* (Burkina Faso), collected the same day. ATQ treatment resulted in a significant reduction in both infection prevalence (GLM (Binomial;Logit): 100 µmol/m^2,^ n=106, df=1, *Χ*^2^=31.997, p<0.0001; 1 mmol/m^2^, n=107, df=1, *Χ*^2^=50.240, p<0.0001) and intensity (GLM (Poisson;Log): 100 µmol/m^2^: n=53, df=1, *Χ*^2^=24.687, p<0.0001, 1 mmol/m^2^: n=59, df=1, *Χ*^2^=31.997, p<0.0001) at both doses, as measured by determination of *P. falciparum* oocyst burden at 7 d pIBM. (**c**) In contrast, transmission of *in vitro-*cultured *P. falciparum* (NF54) is completely blocked when field-derived, lab-adapted insecticide-resistant *An. gambiae* (Bama-R) are exposed to ATQ for 6 min to the maximal effective concentration (EC_99_) for insecticide-susceptible mosquitoes (100 µmol/m^2^) prior to infection. In ATQ-exposed mosquitoes, prevalence was zero, in contrast to the robust infection in mock-exposed controls (Chi^2^, n=160, df=1, *Χ*^2^=93.545, p<0.0001). (**d**) Similarly, transmission of a polyclonal *P. falciparum* West African donor isolate (P5 – *in vitro* culture adapted) was blocked (zero evidence of infection at 7 d pIBM) after ATQ exposure prior an infectious blood meal, despite a strong (prevalence 95.7%, median intensity 59.5 oocysts per midgut) infection in controls (Chi^2^, n=187, df=1, *Χ*^2^=167.995, p<0.0001). Median lines and values are indicated, “n” indicates the number of independent samples. To isolate Oocyst Prevalence and Oocyst Intensity, midgut samples with zero oocysts have been excluded from intensity analysis. Statistical significance is indicated where relevant as follows: ns=not significant, * = p<0.05, **=p<0.01, ***=p<0.001, ****=p<0.0001.

The observation of few parasites surviving exposure may indicate that parasite or mosquito factors in our Burkinabe populations could be reducing the efficacy of ATQ in this assay, potentially including interference from extant insecticide resistance mechanisms in AcVK5 mosquitoes, or reduced ATQ drug sensitivity in *P. falciparum* in this region. To test these possibilities, we initially assayed the efficacy of ATQ against insecticide resistant, Burkina Faso-derived *An. coluzzii* (hereafter Bama-R) with the lab standard *P. falciparum* strain NF54. Reared under pyrethroid selective pressure under otherwise standard laboratory conditions, Bama-R have maintained the parental trait of 100% resistance to permethrin at the WHO discriminating concentration (DC, 696 μmol/m^2^) and exhibit appreciable acute survival at five times this dose (SFig. 1b). Bama-R mosquitoes are segregating for the *kdr* mutation in *para* conferring target site resistance (SFig. 1c), but constitutively overexpress CytochromeP450 genes associated with both metabolic resistance through enhanced small molecule detoxification (SFig. 1d(27)), and cuticular thickening (26) Bama-R females were exposed to the maximal effective concentration for tarsal ATQ (EC_99_, 100 μmol/m^2^, 6 min (25)) or to a vehicle control immediately preceding infection, and parasite prevalence and intensity were determined at 7 d pIBM. While control females were highly infected, with a median of 12 oocysts per infected midgut and 81.25% overall prevalence of infection, no oocysts were observed in females exposed to ATQ, suggesting that insecticide-resistance mechanisms found in highly resistant, natural *Anopheles* populations do not interfere with the transmission blocking activity of ATQ (Fig. 1c).

Next, we established an *in vitro P. falciparum* culture from a polyclonal isolate (P5) collected from a gametocytemic donor from Burkina Faso (28)) and infected the laboratory standard, insecticide susceptible mosquito strain *An. gambiae* (G3). P5 development was 100% suppressed in females treated with the EC_99_ of ATQ (Fig. 1d) such that zero oocysts were observed in midguts, compared to heavy infections—both in terms of infection intensity (median 59.5 oocysts per midgut) and infection prevalence (95.7%)—in controls. Delivery of ATQ to mosquitoes is therefore fully effective against field-derived *P. falciparum* isolates currently circulating in West Africa.

### ATQ prevents the transmission of an artemisinin-resistant *P. falciparum* isolate from the GMS

Given the results obtained with field-derived parasites from Africa, we next tested the ability of ATQ to kill artemisinin resistant parasites from the GMS, where mutations conferring artemisinin resistance occur in a high proportion of *P. falciparum* isolates, constituting a major public health threat. We reasoned that these experiments would also allow us to test the concept of directly targeting drug resistant *P. falciparum* during mosquito development, removing resistance mutations from the parasite population, and thereby “rescuing” ACT efficacy in human treatment. To this end, we used a Cambodian *P. falciparum* patient clone (KH001_029 (5), hereafter ART29) carrying the C580Y mutation in *PfK13* conferring resistance to artemisinin (Fig. 2a). We used the major Asian malaria vector *An. stephensi* for these experiments as initial tests with *An. gambiae* did not produce appreciable infections (SFig. 2). ART29 generated robust infections in control, mock-exposed *An. stephensi*, (median 16 oocysts per midgut, 100% prevalence of infection). Conversely, no oocysts were detected in females exposed to ATQ prior to infection (100 μmol/m^2^, 6 min) (Fig. 2b). These data show that mosquito exposure to antimalarials, such as by incorporation in bed nets, indoor residual sprays (or other contact methods such as eaves tubes (29), could be an effective strategy for reducing the spread of artemisinin resistance both within and between malaria endemic areas, including sub-Saharan Africa.

**Figure 2:**
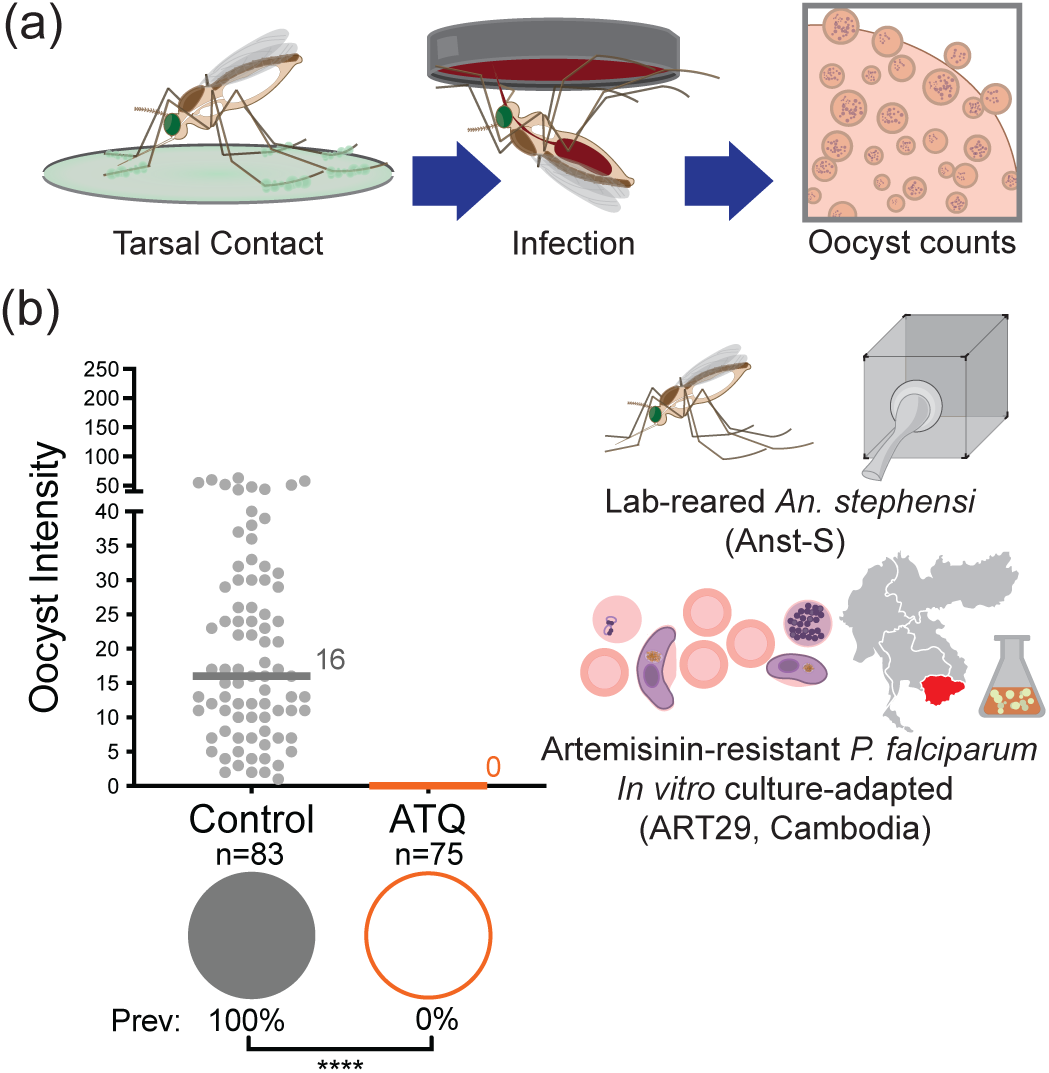
Exposing *An. stephensi* to ATQ blocks an artemisinin-resistant *P. falciparum* patient isolate from Cambodia. **(a)** Experimental scheme. **(b)** Transmission of artemisinin resistant *P. falciparum* (ART29) is completely blocked when *An. stephensi* females (Anst-S) are exposed to ATQ for 6 min to the maximal effective concentration (EC_99_) for insecticide-susceptible mosquitoes. In ATQ exposed mosquitoes, prevalence (indicated by pie charts) was zero despite robust infection in mock-exposed controls (Chi^2^, n=158, df=1, *Χ*^2^=156, p<0.0001). Median lines and values are indicated, “n” indicates the number of independent samples. To isolate Oocyst Prevalence and Oocyst Intensity, midgut samples with zero oocysts have been excluded from intensity analysis. Statistical significance is indicated where relevant as follows: ns=not significant, * = p<0.05, **=p<0.01, ***=p<0.001, ****=p<0.0001.

### ATQ exposure during an ongoing infection delays oocyst growth and decreases sporozoite prevalence

In the field, mosquitoes that contact an antimalarial compound through mosquito-targeted interventions may harbor parasites from a previous blood meal that have already traversed the midgut lumen and formed oocysts. We therefore investigated the effects of ATQ on parasites in which oocyst development is already underway, exposing G3 mosquitoes 6d pIBM (NF54, Fig. 3a). In contrast to females exposed before infection, ATQ had no effect on the prevalence or intensity of infection, as measured at 10d pIBM, suggesting ATQ acts differently on oocysts compared to zygote and ookinetes (Fig. 3b). However, when we measured the size of the developing oocysts, we observed a significant, 45% decrease in the mean oocyst cross-sectional area (Fig. 3c). Oocyst size is a good proxy for rate of growth (30) and as such, when we sampled mosquitoes at a later time point when sporozoite invasion of salivary glands has already occurred (14 d pIBM), we observed a 33% reduction in the prevalence of sporozoites in the salivary glands of ATQ-treated females (Fig. 3d). Similar results were obtained when ATQ exposure instead occurred at 3 d pIBM (SFig. 3). Taken together, these results suggest that ATQ exposure after oocyst formation has a partial cytostatic effect on *P. falciparum*.

**Figure 3:**
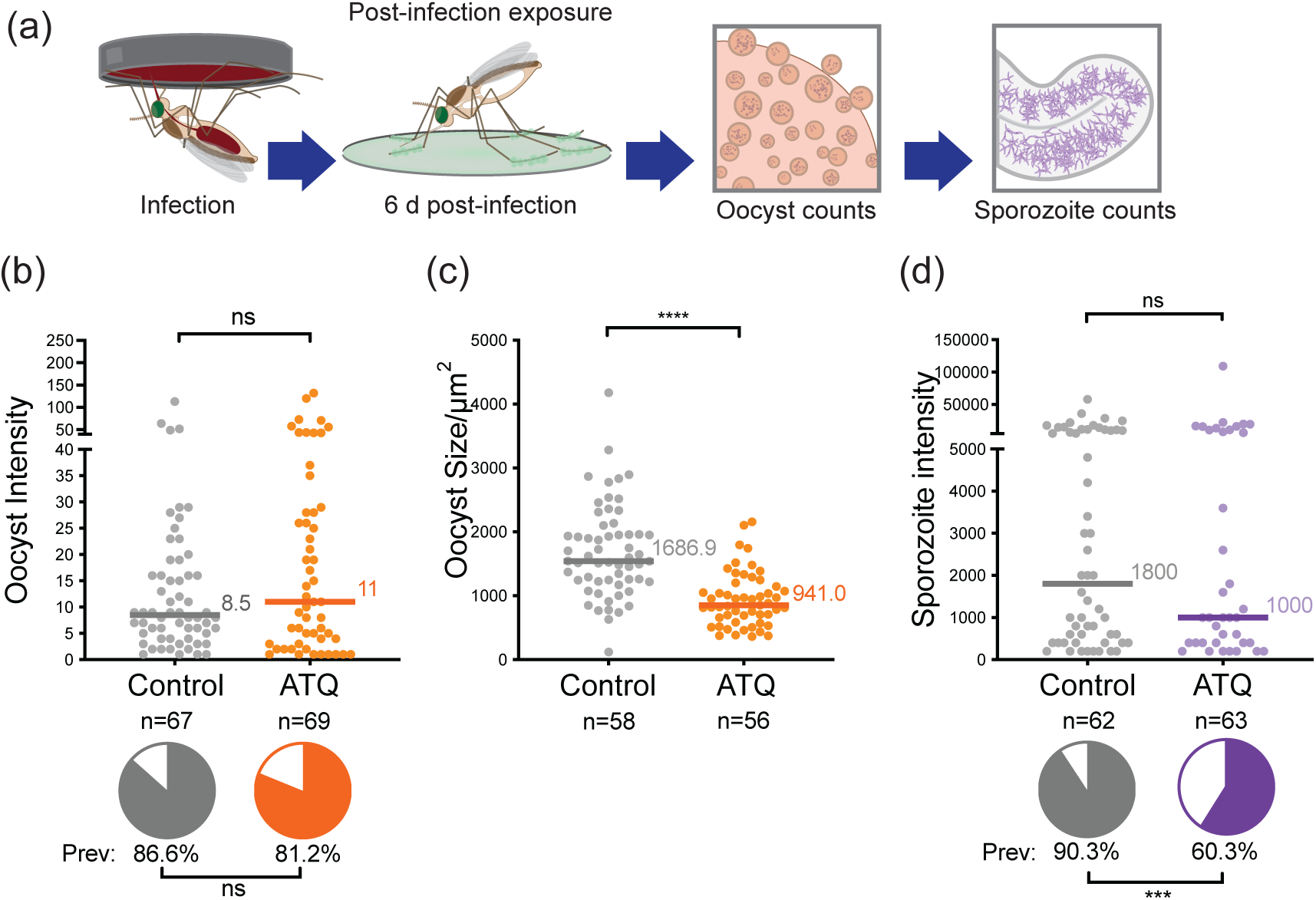
Sporozoite prevalence is significantly reduced after tarsal ATQ exposure during oocyst development when *An. gambiae* (G3) are infected with *P. falciparum* (NF54). (**a**) Experimental scheme. (**b**) There was no effect of 6 d pIBM ATQ exposure on either prevalence (indicated by pie charts) or intensity (indicated by points) of infection determined at 10 d pIBM. Prevalence: Chi^2^, n=136, df=1, *Χ*^2^=0.733, p=0.3919, intensity: Mann-Whitney, n=114, df=1, *U*=1482, p=0.4335. (**c**) ATQ exposure at 6 d pIBM significantly reduced the median cross-sectional area of oocysts relative to control (n=114, df=1, U=531, p<0.0001). (**d**) The prevalence, but not the median intensity of sporozoites in salivary glands was significantly reduced in mosquitoes exposed to ATQ at 6 d pIBM (Chi^2^, n=125, df=1, *Χ*^2^=7.190, p=0.0073). Median lines and values are indicated, “n” indicates the number of independent samples. To isolate Oocyst Prevalence and Oocyst Intensity, midgut samples with zero oocysts have been excluded from intensity analysis. Statistical significance is indicated where relevant as follows: ns=not significant, * = p<0.05, **=p<0.01, ***=p<0.001, ****=p<0.0001.

### Ingestion of an ATQ-glucose solution blocks the establishment of *P. falciparum* infection

To determine whether anti-parasitic compounds in sugar could suppress *Plasmodium* development in the mosquito, we began by testing the efficacy of mosquito ATQ-glucose ingestion against the transmission of *P. falciparum* parasites isolated from gametocytemic donor blood samples collected from gametocyte carriers in Nasso, near Bobo Dioulasso, Burkina Faso. Adult female *An. gambiae,* collected as pupae from larval breeding sites, were denied sugar for 24 h, then given access to an ATQ-treated sugar solution (100 µM ATQ/0.5% v/v DMSO/10% w/v glucose) *ad libitum* for the 24 h preceding infection (Fig. 4a). We observed a striking, 85% reduction in the prevalence of wild *P. falciparum* infection in female mosquitoes that had access to ATQ-glucose prior to infection (Fig. 4b). Importantly, median mosquito survival between control and ATQ treatment groups was not significantly different (SFig. 4), confirming previous findings that atovaquone is not toxic to mosquitoes at parasiticidal concentrations (25). This implies that other *Plasmodium-*specific inhibitors would therefore not impose selective pressures leading to resistance mechanisms in the mosquito.

**Figure 4:**
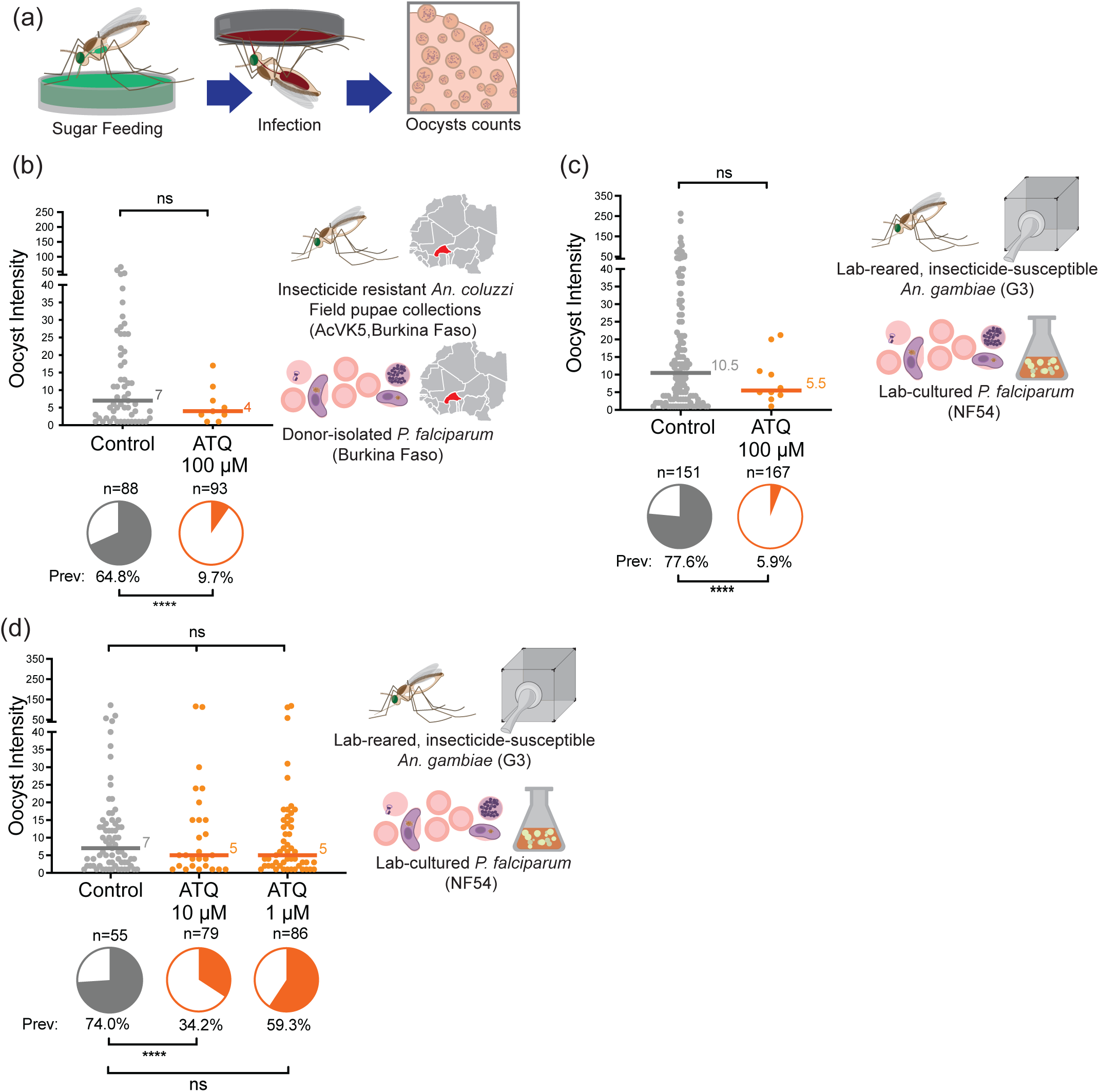
Ingestion of an ATQ-glucose solution prior to *P. falciparum* infection significantly reduces transmission. **(a)** Experimental scheme. **(b)** AcVK5, collected as pupae from field sites in Bama, Burkina Faso, mosquitoes were significantly less likely to become infected with *P. falciparum* (human gametocyte donor sample) after ingestion of 100 µM ATQ/10% w/v glucose (access to treated sugar 24 h prior to infection). The proportion of AcVK5 mosquitoes infected with *P. falciparum* oocysts at 7 d pIBM was reduced from 64.8% in the control group, to 9.7% in mosquitoes given access to ATQ/sugar solution (85.0% reduction, GLM (Binomial;Logit), n=181, df=1, *Χ*^2^= 59.242, p<0.0001). While median intensity of infection in mosquitoes harboring ≥1 oocyst was reduced in ATQ-treated mosquitoes, this reduction was not significant. **(c)** Infection with culture adapted NF54 *P. falciparum* in insecticide susceptible G3 *An. gambiae* produced remarkably similar results. Ingestion of 100 µM ATQ/0.5% DMSO/10% w/v glucose (access to treated sugar 24 h prior to infection) significantly reduced the prevalence of infection at 7 d pIBM (92.4% reduction Chi^2^, n=318, df=1, *Χ*^2^=162.467, p<0.0001) but had no detectable effect on the median intensity of infection (Mann-Whitney, n=128, df=1, *U*=480.5, p=0.3364). **(d)** Significant, dose-dependent inhibition of oocyst prevalence was observed with a 10-fold reduction in ATQ concentration (10 µM: Chi^2^, n=134, df=1, *Χ*^2^=19.275, p<0.0001). A non-significant reduction in prevalence was also observed at 1 µM (Chi^2^, n=141, df=1, *Χ*^2^=2.642, p=0.1041). Median lines and values are indicated, “n” indicates the number of independent samples. To isolate Oocyst Prevalence and Oocyst Intensity, midgut samples with zero oocysts have been excluded from intensity analysis. Statistical significance is indicated where relevant as follows: ns=not significant, * = p<0.05, **=p<0.01, ***=p<0.001, ****=p<0.0001.

Using the same conditions with *in vitro* cultured *P. falciparum* (NF54) and lab-adapted *An. gambiae* (G3) resulted in a remarkably similar infection outcome, with a 92.5% reduction in oocyst prevalence in female *An. gambiae* given access to ATQ-glucose solution relative to controls (Fig. 4c). When ATQ concentration was reduced to 10- and 100-fold ATQ dilutions, we observed progressively reduced, dose dependent effects on prevalence (Fig. 4d).

### ATQ ingestion during an ongoing *P. falciparum* infection impairs sporogony

As mosquitoes may often visit a sugar bait after acquiring an infectious blood meal, we also investigated the impact of ATQ ingestion on ongoing *P. falciparum* (NF54) infections, providing ATQ-treated sugar to G3 females from 2 d pIBM, when ookinetes have escaped the midgut lumen and formed oocysts on the midgut basal lamina (Fig. 5a). This time, as based on our previous results (Fig. 3) we expected a possible cytostatic effect on oocyst growth, we performed a sampling time course to capture oocyst development through mid- to late sporogony (7d, 10d and 14 d pIBM). We also counted salivary gland sporozoites, the end point of parasite development in the mosquito, at 14 d pIBM. In agreement with our previous results, we observed no change in oocyst prevalence and intensity because of ATQ ingestion (SFig. 5). However, we observed an 80.8% decrease in oocyst cross-sectional area relative to controls at 7 d pIBM, which persisted at later time points (89% and 76.3% decreases at 10- and 14 d pIBM, respectively) (Fig. 5b). By 14 d pIBM, ATQ-exposed oocysts had a similar size to 7 d pIBM control oocysts, suggesting a remarkable suppression of growth. Inspection of DAPI-stained infected midguts revealed a stark decrease in the number of nuclear foci, with a single, diffuse DNA signal compared to many condensed foci in oocysts in controls (Fig. 5c). Strikingly, we detected no sporozoites in the salivary glands of mosquitoes given access to ATQ-glucose 14 d pIBM, despit robust infection in controls (Fig. 5d). Combined, these data point to a strong suppression of oocyst growth, DNA replication and sporozoite differentiation after ATQ ingestion, suggesting that besides preventing new mosquito infection, this delivery method would be extremely effective at curbing ongoing infections and transmission.

**Figure 5:**
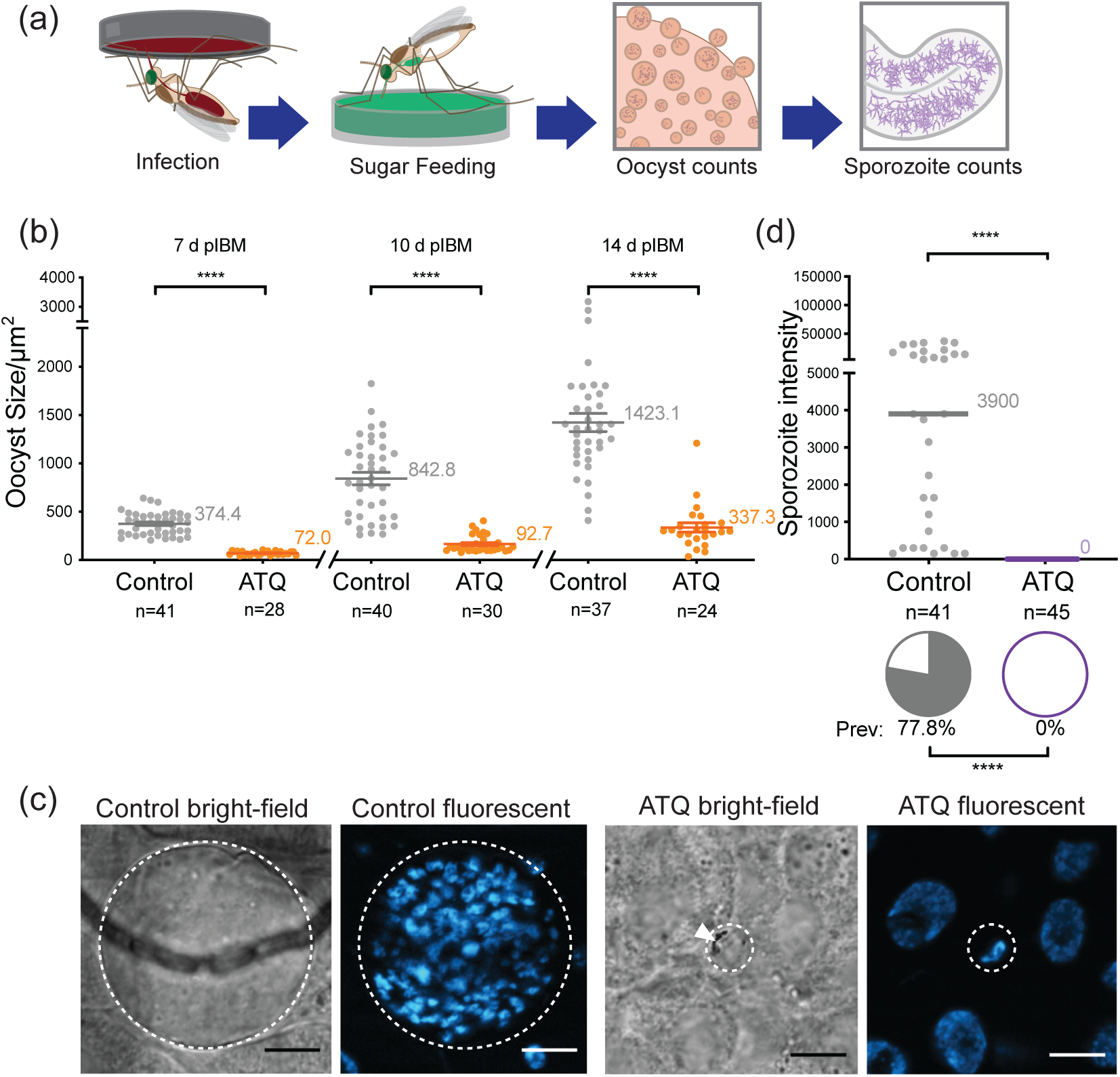
Ingestion of ATQ-glucose after *P. falciparum* (NF54) infection blocks parasite development in *An. gambiae* (G3). **(a)** Experimental scheme. **(b)** Oocyst size over time. Access to ATQ-glucose from 2 d pIBM caused a significant reduction in mean oocyst cross-sectional area (“size”/m^2^) relative to control at 7 d pIBM (2-way T-test, n=69, df=67, *t*=13.63, p<0.0001), 10 d pIBM (2-way T-test, n=70, df=68, *t*=8.998, p<0.001), and 14 d pIBM (2-way T-test, n=61, df=59, *t*=8.793, p<0.0001). Means are indicated, error bars represent the standard error of the mean (SEM). **(c)** Example brightfield, and fluorescent micrographs of control and ATQ-exposed bright-field and DAPI-stained oocysts at 10 d pIBM (outline indicated by dashed line). At this timepoint, the control oocyst is large, and contains many discrete nuclear foci, indicating nuclear division and sporozoite differentiation. In contrast, the ATQ-exposed oocyst is small relative to control and contains a single nuclear focus. Dense hemozoin crystals (white triangle), typically associated with younger oocysts, are visible. Scale bars represent 10 µm, **(d)** Salivary gland sporozoite prevalence and intensity at 14 d pIBM. No sporozoites (0% prevalence, median intensity = 0) were observed in salivary glands samples collected from mosquitoes given access to 100 µM ATQ/0.5% DMSO/10% w/v glucose *ad libitum* (Chi^2^, n=86, df=1, *Χ*^2^=53.202, p<0.0001). Where relevant, median lines and values are indicated, “n” indicates the number of independent samples. To isolate Oocyst/Sporozoite Prevalence and Oocyst/Sporozoite Intensity, midgut samples with zero oocysts have been excluded from intensity analysis. Statistical significance is indicated where relevant as follows: ns=not significant, * = p<0.05, **=p<0.01, ***=p<0.001, ****=p<0.0001.

## Discussion

In this study we demonstrate the strong potential of incorporating antimalarials into LLINs, IRS and ATSBs to stop transmission of endemic parasites by insecticide-resistant *Anopheles*. Our proof of principle compound ATQ was very effective at killing a polyclonal, field-derived parasite isolate from sub-Saharan Africa, showing that an antimalarial-based LLIN or IRS could suppress transmission even in areas of malaria hyperendemicity, where vector pyrethroid resistance is exceptionally high, and multiple parasite haplotypes coexists (31, 32).

Tarsal ATQ exposure against wild *An. coluzzii* (AcVK5) collected as pupae from breeding sites was able to strongly suppress the transmission of *P. falciparum* isolates circulating in children. Interestingly, in these infections, carried out in Burkina Faso, we observed a small number of “breakthrough” oocysts at ATQ doses that are non-permissive in tests using, respectively, lab-adapted and susceptible NF54 parasites and G3 mosquitoes (25). We reasoned that this marginal reduction in efficacy could be due to extant insecticide resistance circulating in wild *Anopheles gambiae* s.l populations in Burkina Faso, including the wild AcVK5 mosquitoes used for these experiments. While clearly target-site resistance mechanisms are unlikely to impact the activity of antimalarials like ATQ, both cuticular and metabolic resistance could potentially interfere with function by limiting compound uptake and stability, respectively. The CYP450 monooxygenases associated with pyrethroid metabolic detoxification in *An. gambiae* s.l. have been shown to confer some degree of resistance to a structurally and functionally diverse array of compounds including the insecticide DDT, the juvenile hormone agonist pyriproxyfen (33), and several arthropod mitochondrial complex I inhibitors including the otherwise promising insecticide fenpyroximate (34). Similarly, cuticular resistance, where the waxy exocuticle of the mosquito has thickened as an adaptation to insecticide pressure (11), could slow or eliminate uptake of other small molecules. However, tarsal ATQ (at the EC_99_, 100 µmol/m^2^ for 6 min) completely abrogated parasite infection in Bama-R mosquitoes, which are derived from the parental AcVK5 population and exhibit a similar, high level of insecticide resistance combining the additive effects of metabolic and cuticular resistance components (SFig. 1, (26)). This observation suggests that mosquito pyrethroid resistance status, or other vector factors, did not affect tarsal antimalarial efficacy in our experiments.

Similarly, in the reciprocal experiment, transmission of the culture-adapted Burkinabe *P. falciparum* isolate P5 was blocked in an insecticide susceptible *An. gambiae* lab strain (G3), again indicating that parasite factors are likely not responsible for the observed reduction in efficacy. Clinically-induced resistance to ATQ in *P. falciparum* is associated with mutation at position 268 of the mitochondrial gene *cytochrome B* (35), while *in vitro* selection typically results in mutation elsewhere in the gene (36–38). ATQ is not in clinical use in sub-Saharan Africa, so it is unlikely that ATQ resistance-conferring mutations are circulating in Burkina Faso at any appreciable frequency. However, while certain ATQ-resistance mutations induce a transmission defect in both *P. berghei* and *P. falciparum -* resulting in failure to establish infection in mosquitoes (37) there are contradictory findings in the literature (38), and it has been shown that naturally occurring ATQ resistance-conferring mutations can persist at ultra-low frequency in parasite populations through mitochondrial heteroplasmy (39). Thus while the Burkinabe isolate (P5) used in our lab-based studies was susceptible to ATQ *in vitro*, and Sanger sequencing of *Cytochrome B* from both P5 asexual blood stages and oocysts showed that these parasites are wild-type, we cannot rule out the possibility that either cryptic parasite factors, lost during the short P5 laboratory adaptation period, or specific vector-parasite interactions in wild populations are responsible for the observed reduction in ATQ efficacy in our Burkina-based experiments. The observed decrease in *P. falciparum* numbers remains extremely significant (above 96% total inhibition at 100 µmol/m^2^). Taken together, these data demonstrate that mosquito-targeted antimalarial exposure can bypass currently circulating insecticide resistance mechanisms, maintaining activity even in areas where conventional insecticides have ceased to be effective — an essential trait of any new mosquito-targeted tool.

ATQ exposure also killed artemisinin-resistant *P. falciparum* parasite ART29 in *An. stephensi*. *De novo PfK13* mutations associated with *in vitro* resistance, already highly prevalent in Cambodia and other part of the GMS, have now been detected in Uganda, Tanzania and Rwanda (14–19) and the recent invasion and spread of *An. stephensi* to the horn of Africa (20) may facilitate the spread of these parasites. Widespread artemisinin resistance in Africa, a region where malaria prevention is already challenging due to insecticide resistance, would be a major public health concern. Although our results were somewhat expected given ART29 parasites are ATQ-sensitive *in vitro* during asexual development (40), they are an important proof of concept, both that drug-based mosquito-targeted interventions could be developed to specifically contain and eliminate parasite haplotypes conferring resistance to human antimalarial therapeutics, and that tarsal uptake of antimalarials function similarly in *An. stephensi*. Antimalarial pressure directed at different drug targets in the human and mosquito life stages could effectively suppress the spread of resistance mutations selected in either host and to either target, allowing the possibility for close integration of human- and mosquito-targeted antimalarial interventions in the future. Importantly, mosquito-targeted antimalarials attack the parasite during an extreme population bottleneck and in a non-cyclical stage in its life cycle (41–43), reducing the probability of selection of both extant and *de novo* resistance mutations during mosquito-stage drug challenge compared to treatment during the human asexual cycle. Nevertheless, even the possibility for selection of resistance during parasite sporogonic development means, as a fundamental principle, any mosquito-targeted antimalarial compound integrated into LLIN or IRS must not share a mode of action with current human antimalarial therapeutics. Thus, despite its efficacy in our studies, ATQ could not be responsibly incorporated into a mosquito-targeted intervention due to its use as a human prophylactic and therapeutic drug (44), making identification of additional active compounds in diverse mode of action classes a priority.

Ingestion of sugar solutions containing ATQ before *P. falciparum* infection blocked transmission in both field-derived and *in vitro* cultured *P. falciparum*. Such close agreement between these experiments, carried out at different sites with different parasites and mosquitoes, is a testament to the promise of this strategy, demonstrating its high effectiveness in spite of the inherent variability of mosquito sugar feeding behavior (45). Furthermore, continued mosquito access to ATQ-sugar post infection reduced oocyst growth resulting in absence of salivary gland sporozoites. These results support the use of antimalarials in ATSB-like strategies to reduce residual malaria transmission carried out by mosquitoes that predominantly rest and feed outside and thereby avoid both LLINs and IRS. While current proposed ATSB designs rely on insecticidal ingredients (23, 24, 46), the use of antimalarials in sugar baits could act as a more specific and environmentally benign paradigm for this promising intervention should suitable antimalarial ingredients be identified (47–49). ATQ uptake via tarsal contact at 3- or 6-days post infection also significantly reduced oocyst growth, resulting in an appreciable reduction in the proportion of salivary gland sporozoite-positive mosquitoes. *P. falciparum* is therefore also vulnerable to inhibition through tarsal exposure during the oocyst stage, which is by far the longest developmental stage in the mosquito, taking between 7-10 days depending on the frequency of blood feeding (30), and thus is the parasite life stage most likely to encounter a mosquito-targeted intervention. As such, the ability to stall or kill oocysts and sporozoites is a highly desirable quality for mosquito-targeted antimalarials.

ATQ targets the ubiquinol oxidation (Qo) site of cytochrome b (50), a key element of the mitochondrial electron transport chain (mtETC) which has dual roles in both ATP generation through oxidative phosphorylation and DNA replication through ubiquinone-mediated redox of dihydroorotate dehydrogenase (51). Thus, inhibition of either DNA replication or ATP production, or both, could explain the cytostatic effect observed here. Consistent with our findings, previous studies have shown that disruption of ATP production through knock-out of components of the tricarboxylic acid cycle in *P. falciparum* (52) and mtETC in *P. berghei* (53–55) can cause oocyst arrest. Moreover, chemical inhibition of *P. vivax* DNA replication during sporogony was also sporontocidal (56), suggesting mitochondrial inhibitors could also be utilized to prevent transmission of these widespread human malaria parasites. Identifying the specific mechanism by which ATQ and other mtETC inhibitors affect sporogonic development in *Plasmodium* is an interesting area for further study.

Although at an early stage, mosquito-targeted antimalarials have the potential to be an effective element to drive malaria incidence down. To this end, identifying more compounds with strong antiparasitic activity during the mosquito stages of *P. falciparum* development – and in particular compounds with sporogony-specific activity – will be a crucial next step, and one that should leverage the extensive libraries of known antimalarials. Indeed, one of the key strengths of this approach is the potential to exploit and repurpose compounds that are otherwise unsuitable for human therapeutic use, whether due to toxicity, poor bioavailability, poor kinetics, or other limiting factors. By integrating human and mosquito-based interventions, this strategy will extend and protect the efficacy of human therapeutics and vector control strategies, giving malaria control efforts renewed vigor.

## Materials and Methods

### Mosquito lines, insecticide resistance selection and husbandry

*Anopheles* spp. mosquito populations used in this study were: 1) wild *An. coluzzii* captured as pupae from breeding sites, described below. 2) Laboratory-reared *An. gambiae* obtained from an outbred colony established in 2019 and repeatedly replenished with F1 from wild-caught females collected in Soumousso (11°23′14′′N, 4°24′42′′W).3) *An. gambiae* G3 (“G3”), a highly lab-adapted, insecticide-susceptible strain competent for *P. falciparum* of African origin. 4) *An. stephensi* (Anst-S), a similarly lab-adapted, insecticide-susceptible strain competent for *P. falciparum* of both African and Asian origin received as a gift from The Institute of Molecular Medicine, University of Lisbon, Portugal. 5) *An. coluzzii.* Bama-R (“Bama-R”) a colony established through hybridization of the F1 progeny of female *An. coluzzii* collected from Vallee du Kou, Burkina Faso, with our G3 colony. Since establishment, Bama-R has been kept under frequent permethrin selection pressure and exhibits a consistent pyrethroid resistance phenotype. At the time of this study, (F17-20) Bama-R females where highly resistant to pyrethroids, exhibiting 97% survival in standard WHO insecticide resistance assays — briefly, 1 h exposure to the WHO discriminating concentration (DC, 275 mg/m^2^) of permethrin-impregnated papers with mortality scored at 24 h post-exposure — and 43% survival at 5x the DC. Except for selection of resistance in Bama-R, all mosquito colonies were maintained identically at 26 ⁰C ± 2 ⁰C and 80% ± 10%. relative humidity (RH). Larvae were cultured in 2-liter (l) catering pans in 500 ml distilled water (dH_2_O) under an optimized density and feeding regimen. At the onset of pupation, pupae were separated from larvae using a vacuum aspirator, collected in dH_2_O, and placed in a 30×30×30 cm cage (Bugdorm, Megaview Science Co, Ltd, Thailand). After emergence, adult mosquitoes had access to separate sources of 10% glucose (Sigma Aldrich, US) and dH_2_O *ad libitum*. For colony maintenance, 5–7-day-old adults were provided a blood meal of donated human blood using an artificial membrane feeding system (Hemotek, UK). For mosquito colony 2) females were maintained on rabbit blood by direct feeding (protocol approved by the national committee of Burkina Faso; IRB registration #00004738 and FWA 00007038) and adult males and females fed with a 5% glucose solution. Larvae were reared at a density of about 300 first-instar larvae in 700 ml of water in plastic trays and fed with Tetramin Baby Fish Food (Tetrawerke, Melle, Germany).

### P. falciparum strains and culture

*P. falciparum* strains used in this study were: 1) *P. falciparum* NF54. NF54 is the drug-susceptible standard strain for mosquito transmission studies, obtained from BEI Resources in 2014. This parasite culture was received through a material transfer agreement (MTA) with the laboratory of Dr. Carolina Barillas-Mury. 2) *P. falciparum* P5. P5 is polyclonal (n=3, KMMM, KMKM, RMMM), as determined by MSP1 PCR genotyping following standard procedures (28), and has been culture-adapted from a blood sample contributed by a gametocytemic malaria donor in Burkina Faso in 2017. 3) *P. falciparum* KH001_029 “ART29”. ART29 is a *P. falciparum* monogenomic parasite isolate obtained from an infected human patient in Pursat, Cambodia (KH1 clade) between 2011 and 2013 as part of the TRAC I initiative (5). This parasite carries the *PfK13* mutation C580Y associated with resistance to artemisinin. ART29 has clear phenotypic artemisinin resistance, both as determined by *in vivo* clearance time (11.8 h) and *in vitro* ring-stage survival (22.6%) (57), alongside resistance to other antimalarial drugs, including mefloquine and chloroquine, but is not resistant to piperaquine or ATQ (40). These parasites generate robust infections in *An. stephensi* mosquitoes, but not *An. gambiae* (SFig. 2).

For mosquito infection with gametocytemic donor blood samples, *P. falciparum* samples were collected and prepared for infection as described previously (58). Briefly, *P. falciparum* gametocyte-positive whole blood samples were collected from 5–13-year-old donors from the villages surrounding Bobo Dioulasso, Burkina Faso as part of a separate study. Red blood cells were isolated by centrifugation of a 4 ml aliquot of donor blood followed by resuspension in *Plasmodium* naïve human AB serum. Blood samples were then provided to mosquitoes using a custom blown, water heated glass feeder. For mosquito infection with *in vitro* cultured parasites, females were transferred to a secure malaria infection facility and provided a 14-21 d post-induction stage V *P. falciparum* gametocyte culture using a custom blown, water heated glass feeder. For all infection experiments, within 24 h of infection, partially engorged or unfed mosquitoes were collected by vacuum aspiration and discarded. To determine oocyst burden, between 7 and 14 d pIBM, infected mosquitoes were collected by vacuum aspiration, and dissected to isolate the midgut. The oocyst burden was determined after staining midguts with 0.2% w/v mercurochrome and examination under a 40x air objective inverted compound light microscope (Olympus, US). For sporozoites, at 14 d pIBM infected mosquitoes were collected by vacuum aspiration and beheaded. The mosquito salivary glands were extracted into RPMI media by applying pressure to the lateral thorax. *P. falciparum* sporozoites were isolated by homogenization and centrifugation of salivary gland material, followed by resuspension in a known volume of RPMI. Sporozoites were counted using a disposable haemocytometer under a 20x air objective inverted compound light microscope (Olympus, US). All *P. falciparum* strains were cultured and induced to form gametocytes using standard protocols (59, 60). All strains have been confirmed to be *P. falciparum* by PCR followed by DNA sequencing of the amplified products (61) and have been confirmed free of mycoplasma infection.

### Pre- and post-infection tarsal contact infection assays

Tarsal exposure plates were prepared as described previously (25). Briefly, for 100 µmol/m^2^ plate concentrations, 0.1 ml of this solution of a 0.1% w/v solution of ATQ in acetone was diluted with 1 ml additional acetone and spread onto a 6 cm diameter (0.002628 m^2^) glass petri dish. For 1 mmol/m^2^, 1 ml of 0.1% w/v ATQ/acetone solution was added directly to the plate. Plates dried for a minimum of 4 h, with agitation on a lateral shaker at room temperature. For pre-infection exposure, 30 min prior to infection, 3-5 d old virgin female mosquitoes were incubated on either ATQ-coated plates, or an acetone treated control, for 6 min. To prevent crowding and agitation, a maximum of 25 mosquitoes were exposed per plate, with all exposures occurring in parallel. For post-infection exposure, due to biosafety considerations, compound exposures were carried out in serial, with a maximum of 10 infected mosquitoes per plate.

### Pre- and post-infection sugar feeding infection assays

For experiments involving compound-treated sugar solutions, 20 mM stock solutions of each active ingredient (AI) were prepared in 100% DMSO. Each 20 mM stock solution was diluted 200-fold in 10% w/v glucose to achieve the final working concentration of 100 µM AI/0.5% DMSO/10% w/v glucose. For pre-infection sugar feeding experiments, 2-4 d post-emergence females were denied access to glucose or water for 24 h and then provided access to either the test solution, or a control solution of 0.5% v/v DMSO/10% glucose, for 24 h. After this time had elapsed, all mosquitoes were provided with an infectious *P. falciparum* blood meal as described above. Infected mosquitoes were provided with untreated 10% w/v glucose *ad libitum* for the remainder of the experiment. For post-infection sugar-feeding experiments, 3-5 d post-emergence female mosquitoes were infected with *P. falciparum* and denied access to glucose or water for 48 h pIBM. After this time, infected mosquitoes were continuously provided either 100 µM ATQ/0.5% DMSO/10% w/v glucose or 0.5% DMSO/10% w/v glucose as control, *ad libitum.* Sugar feeders were replaced every 48 h for the remainder of the experiment, up to 14 d pIBM.

### Statistical analyses

Statistical analyses were carried out using GraphPad Prism v8.4.2 for MacOSX (GraphPad Software Inc., USA) and JMP Pro 15 (SAS Corp. US).

### Data generated from donor isolated gametocytes

For infections with donor isolated gametocytes (Fig. 1(b), Fig. 4(b)), prevalence and intensity of infection were analyzed using more complex statistics to account for between-replicate effects of different human gametocyte donors. **For prevalence:** we constructed a General Linear Model as follows, independent variable/y “Infected?” (Two-level, categorical (yes/no)), with cofactors “Treatment” (Two-level, categorical (Control/ATQ)) and “Gametocyte Donor” (Two-level, categorical (Donor 1/Donor 2)), we also included the interaction term Treatment*Gametocyte Donor to detect higher level effects. As the output was categorical, the GLM model was run with a binomial distribution and logit link-function. To achieve the best model fit, we iteratively removed cofactors from the model, and selected the model output with the lowest corrected Akaike information criterion (AICc). In all cases, the best model fit included both cofactors, but excluded the interaction term. **For intensity:** Independent variable “Oocyst Count” (Continuous, positive integer) with cofactors “Treatment” (Two-level, categorical (Control/ATQ)) and “Gametocyte Donor” (Two-level, categorical (Donor 1/Donor 2)), and the interaction term Treatment*Gametocyte Donor. To account for the overdispersion typical of parasite count data, we again took an iterative approach to model construction. Data were analyzed using both a GLM using a Poisson distribution, (link function: log; overdispersion parameter estimated by Pearson Chi-square/DF) and Generalized Regression with a Negative Binomial distribution. Relative model quality was determined by comparison of AICc for each distribution function (For GLM, with and without the overdispersion correction) and by iterative removal of cofactors. The highest quality model fit was an overdispersion-corrected Poisson/Log GLM with both cofactors but without the interaction term.

### Data generated from in vitro experiments

For all other infections, differences in prevalence were analyzed by Chi^2^. In experiments where both treatment groups had individuals that produced >0 oocysts, differences in median oocyst burden between groups (intensity of infection) was analyzed using a Mann-Whitney Mean Ranks test. For multiple comparisons, differences in prevalence between multiple groups were determined using pairwise Chi^2^ corrected for multiple comparisons (Bonferroni). Similarly, multiple comparisons of intensity were carried out using a Kruskal-Wallis test with Dunn’s *post hoc*.

## Author Contributions

DGP and FC wrote the manuscript. DGP, TL and FC designed the experiments. DGP and TL carried out statistical analysis. DGP, ASP, EM, DFSH, PP, RSY, TL and NS carried out infection experiments. DFSH, RSY, TL, WRS, SKV, and SB collected, established, maintained, and assayed parasite lines. KLA established and reared insecticide-resistant mosquito lines and carried out resistance assays. TL, AD, RKD, DFW, and FC, provided funding and oversight. All authors read and approved the manuscript prior to submission.

The authors state that they have no conflicting interests.

---

**Supplementary Figure 1:**
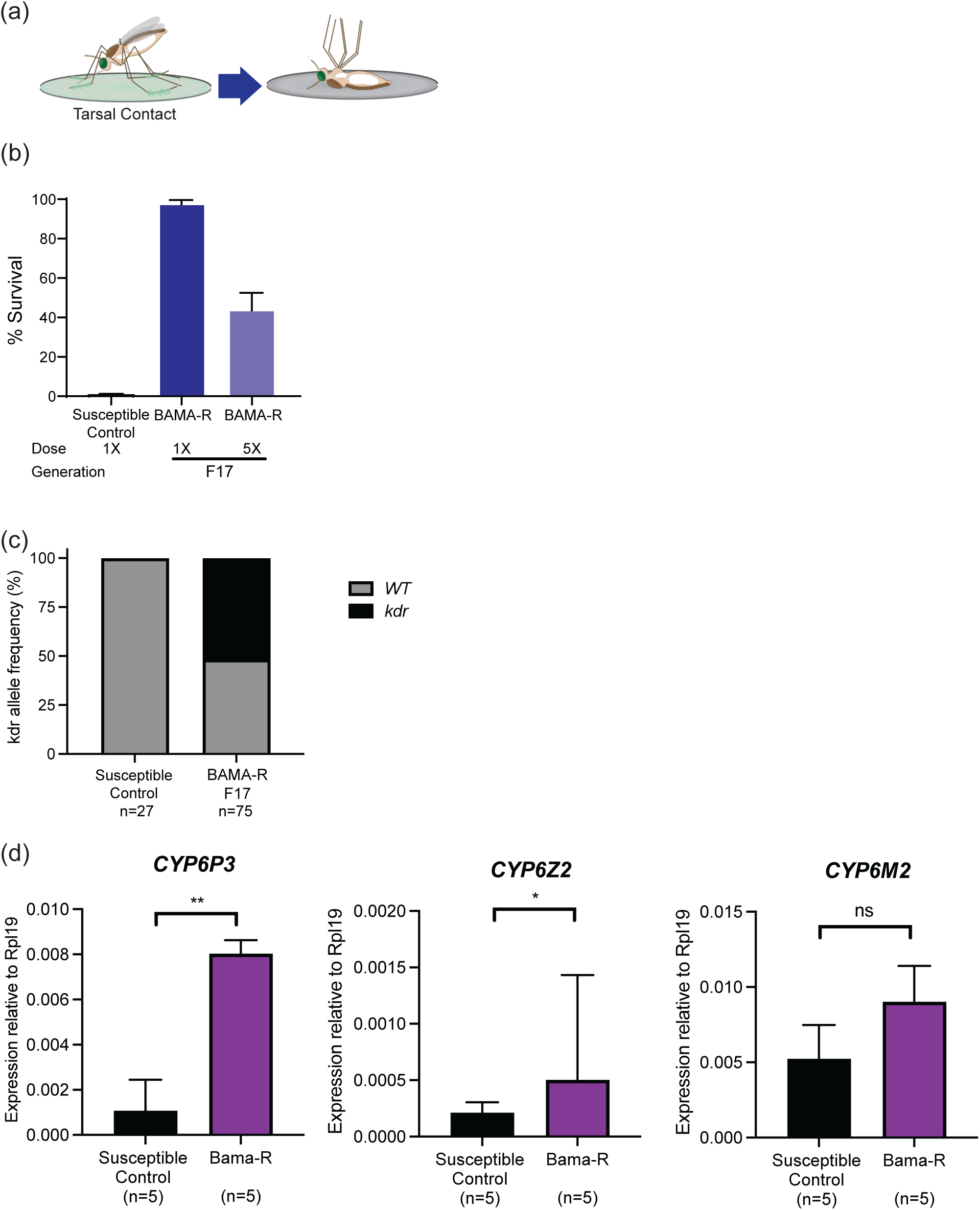
*An. gambiae* Bama-R mosquitoes are highly resistant to permethrin. **(a)** Experimental scheme. **(b)** G3 females were 0% resistant at the DC, while 97% of exposed Bama-R females survived at the same dose, and 43% survived exposure to 5xDC, indicating extreme permethrin resistance. Mean survival ± SEM from 3 replicates is indicated. **(c)** Bama-R mosquitoes are segregating for the *kdr* allele conferring target site resistance to pyrethroids. Allele frequency at generation F17 was 51.7% (n=75), indicating that the observed resistance phenotype is the result of additional modalities. **(d)** qPCR analysis shows that key cytochrome P450 genes associated with metabolic insecticide resistance are constitutively upregulated in Bama-R females compared to a susceptible control (G3). Median expression level normalized to the *Anopheles* housekeeping gene *rpl19* are shown, error bars represent the interquartile range (IQR). Expression levels for *Cyp6P3* and *Cyp6Z2* were significantly elevated compared to a phenotypically susceptible control (*Cyp6P3*, Mann-Whitney, n=10, df=1, *U=*0, p=0.0079; *Cyp6Z2*, Mann-Whitney, n=10, df=1, *U=*2, p=0.0317) while *Cyp6M2* was not significantly upregulated. Statistical significance is indicated where relevant as follows: ns=not significant, * = p<0.05, **=p<0.01, ***=p<0.001, ****=p<0.0001.

**Supplementary Figure 2:**
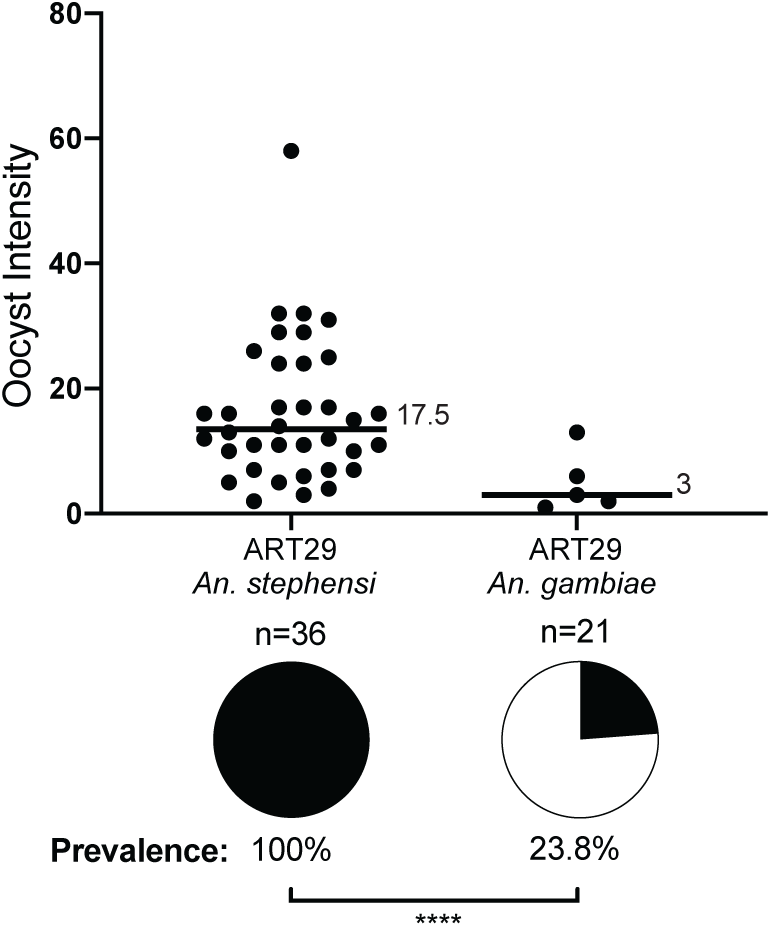
Development of the artemisinin-resistant *P. falciparum* parasite ART29 in *An. stephensi* and *An. gambiae*. Female G3 (*An. gambiae*) and Anst (*An. stephensi*) were provided with a blood meal containing mature ART29 gametocytes. Outcome of infection was determined at 7 d pIBM by oocyst count. While ART29 exhibited poor infectivity in *An. gambiae* (23.8% infection prevalence), they established robust infections in *An. stephensi* (100% infection prevalence).

**Supplementary Figure 3:**
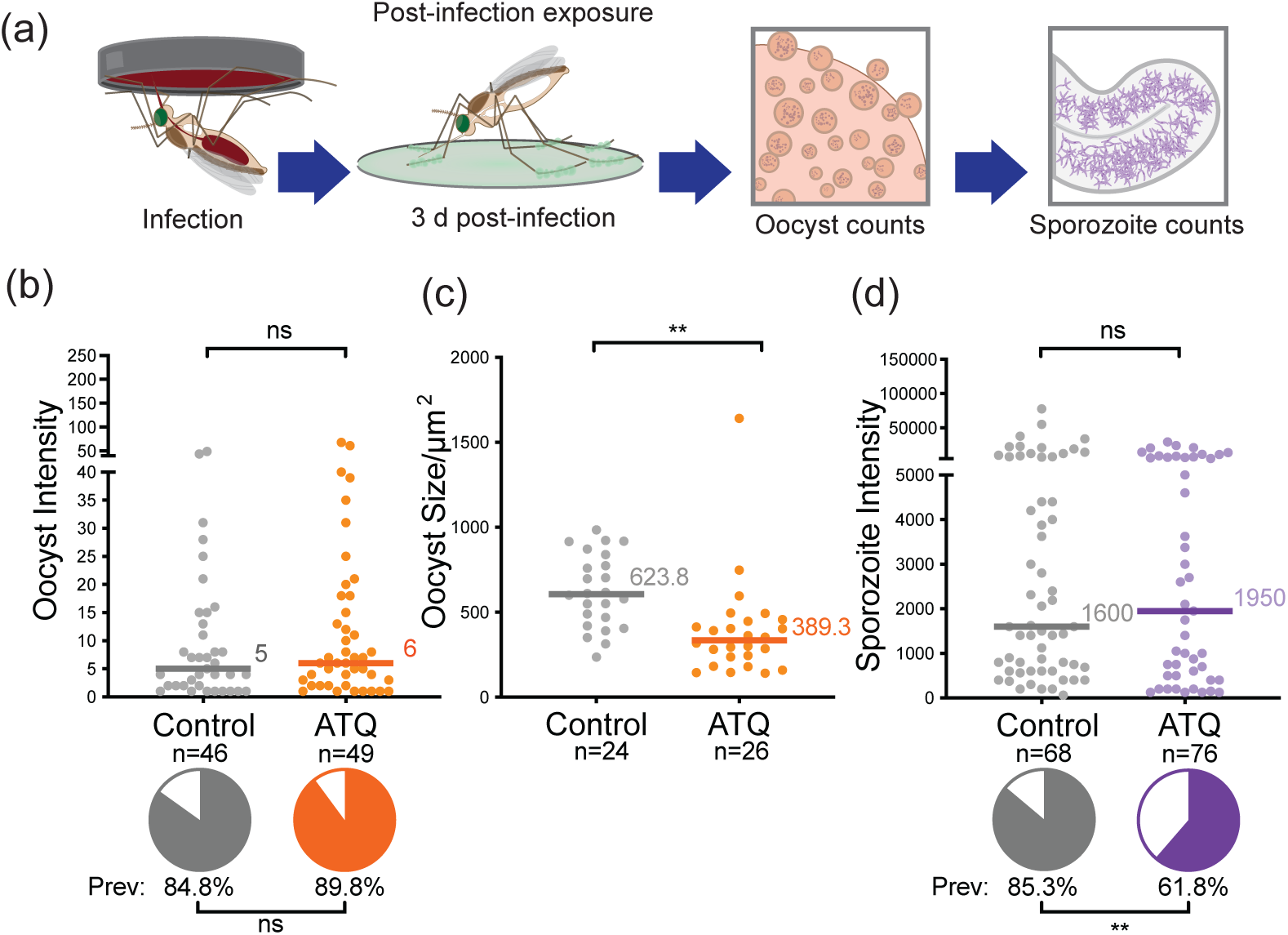
The proportion of *An. gambiae* mosquitoes with salivary gland sporozoites is significantly reduced after tarsal ATQ exposure at 3 d pIBM. **(a)** Experimental scheme. **(b)** There was no effect of 3 d pIBM ATQ exposure on either prevalence (indicated by pie charts) or intensity (indicated by points) of infection determined at 10 d pIBM. Prevalence: Chi^2^, n=95, df=1, *Χ*^2^=0.540, p=0.4623, intensity: Mann-Whitney, n=80, df=1, U=717.5, p=0.4530. **(c)** ATQ exposure at 3 d pIBM significantly reduced the median cross-sectional area of oocysts at 7 d pIBM relative to control (Chi^2^, n=50, df=1, *U*=110, p<0.0001). **(d)** The prevalence, but not the median intensity of *P. falciparum* sporozoites in mosquito salivary glands was significantly reduced in mosquitoes exposed to ATQ at 3 d pIBM (Chi^2^, n=144, df=1, Χ^2^=9.995, p=0.0016). Median lines and values are indicated, “n” indicates the number of independent samples. To isolate Oocyst/Sporozoite Prevalence and Oocyst/Sporozoite Intensity, midgut samples with zero oocysts have been excluded from intensity analysis. Statistical significance is indicated where relevant as follows: ns=not significant, * = p<0.05, **=p<0.01, ***=p<0.001, ****=p<0.0001.

**Supplementary Figure 4:**
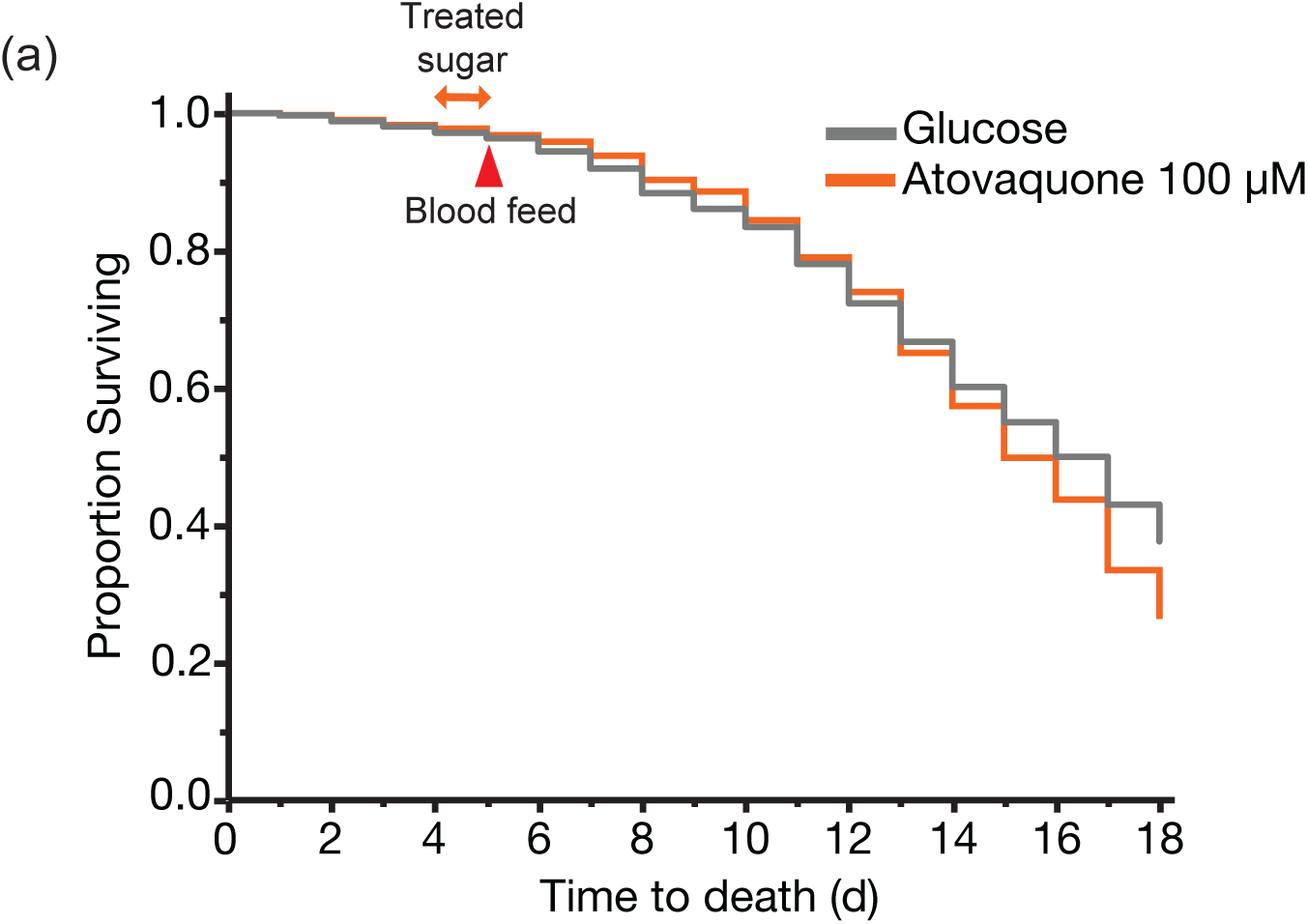
AcVK5 survival and infectivity after ingestion of ATQ/glucose. **(a)** Survival prior to- and following 24 h access to 100 µM/10% w/v Glucose/0.5 v/v/ DMSO (4 d post emergence, orange arrow), followed by *P. falciparum* (donor blood) infection (5 d post emergence, red arrow). Ingestion of ATQ/glucose had no impact on the survival of AcVK5 mosquitoes relative to a control group provided with 10% w/v glucose/0.5% v/v DMSO (Log-Rank Survival, n=1374, df=1, Χ^2^=1.3795, *p*=0.2402).

**Supplementary Figure 5:**
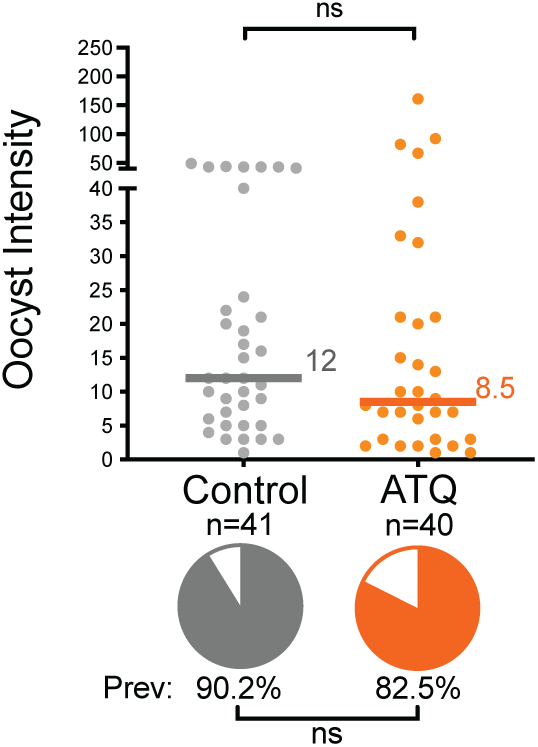
Ingestion of an ATQ-glucose solution after *P. falciparum* infection has no effect on oocyst prevalence or intensity. There was no difference relative to controls in either intensity (n=68, df=1, *U*=495.5, p=0.3257) or prevalence (n=81, df=1, *Χ*^2^=2.441, p=0.1182) of oocysts at 10 d pIBM in females with continued access to 100 µM ATQ/0.5% DMSO/10% w/v glucose from 2 d pIBM - 14 d pIBM. Median lines and values are indicated, “n” indicates the number of independent samples. To isolate Oocyst Prevalence and Oocyst Intensity, midgut samples with zero oocysts have been excluded from intensity analysis. Statistical significance is indicated where relevant as follows: ns=not significant, * = p<0.05, **=p<0.01, ***=p<0.001, ****=p<0.0001.

